# Sequence-encoded Spatiotemporal Dependence of Viscoelasticity of Protein Condensates Using Computational Microrheology

**DOI:** 10.1101/2024.08.13.607792

**Authors:** Dinesh Sundaravadivelu Devarajan, Jeetain Mittal

**Author notes:** Corresponding authors’.

## Abstract

Many biomolecular condensates act as viscoelastic complex fluids with distinct cellular functions. Deciphering the viscoelastic behavior of biomolecular condensates can provide insights into their spatiotemporal organization and physiological roles within cells. Though there is significant interest in defining the role of condensate dynamics and rheology in physiological functions, the quantification of their time-dependent viscoelastic properties is limited and mostly done through experimental rheological methods. Here, we demonstrate that a computational passive probe microrheology technique, coupled with continuum mechanics, can accurately characterize the linear viscoelasticity of condensates formed by intrinsically disordered proteins (IDPs). Using a transferable coarse-grained protein model, we first provide a physical basis for choosing optimal values that define the attributes of the probe particle, namely its size and interaction strength with the residues in an IDP chain. We show that the technique captures the sequence-dependent viscoelasticity of heteropolymeric IDPs that differ either in sequence charge patterning or sequence hydrophobicity. We also illustrate the technique’s potential in quantifying the spatial dependence of viscoelasticity in heterogeneous IDP condensates. The computational microrheology technique has important implications for investigating the time-dependent rheology of complex biomolecular architectures, resulting in the sequence-rheology-function relationship for condensates.

## Introduction

Characterizing the spatiotemporal evolution of biomolecular condensates is crucial for understanding their role in modulating cellular biochemistry^1-3^ and how they transform into pathological aggregates.^4-6^ Liquid-liquid phase separation through multivalent interactions in a protein sequence can drive the formation of these condensates.^7-12^ Liquid-like (viscous) behavior of these cellular compartments is thought to define their functional landscape, by enabling extreme dynamics^13^ and efficient transport of biomolecules that can aid in biochemical processes.^14^ However, recent investigations have demonstrated the loss of condensate liquidity over time, yielding dominant elastic behavior that may eventually promote the formation of solid fibrillar states.^15-17^ Thus, it is critical to accurately quantify the viscoelastic spectrum of biomolecular condensates to establish how the protein sequence governs their (dys)functional paradigm.^18^

Transitions in the material states of protein condensates can occur due to a multitude of factors, e.g., post-translational modifications that alter sequence charge patterning or mutations that alter sequence hydrophobicity.^19-21^ Such sequence alterations commonly occur in intrinsically disordered proteins (IDPs) or regions (IDRs), which are prevalent in biocondensates, thereby causing a speed-up or slowdown in their dynamics. Deciphering the sequence-encoded molecular interactions of IDPs that dictate prominent dynamical changes in conjunction with the measurements of viscoelasticity are essential for establishing the sequence-rheology-function relationship of condensate biology. Simulations can serve as a computational lens into the molecular interactions of condensates and a tool for accurately measuring their rheological properties. For example, the viscosity and viscoelasticity of IDP condensates can be quantified using equilibrium molecular dynamics (MD) simulations along with the Green-Kubo (GK) relation and nonequilibrium MD simulations.^22-26^ While these techniques are useful for measuring the bulk rheology of single-component systems, they suffer from the inability to capture the spatial variations in viscoelasticity seen in multi-component in vitro and in vivo condensates. Knowledge of the spatial dependence of viscoelastic properties within the condensates can provide important insights for studying how the partitioning and transport of small drug molecules into condensates depends on their local environment.^27^

Particle tracking experimental microrheology is widely used for investigating the time-dependent bulk viscoelasticity and viscosity of in vitro phase-separated droplets displaying different material characteristics ranging from liquid-like to solid-like behaviors.^28-33^ It is a highly sought-after method because it requires only small volumes of biological samples and enables a label-free approach where the biomolecules need not be fluorescently tagged.^34, 35^ In addition, the technique has the potential to quantify the local viscoelasticity of heterogeneous condensate systems.^34^ The technique relies on connecting the probe particle motion in a complex fluid system and its microscopic viscoelastic properties through continuum mechanics.^36, 37^ A computational analogue of the technique via MD simulations has been shown to yield quantitatively accurate viscoelastic modulus for the homopolymer and colloidal systems,^38-41^ but a rigorous implementation of it remains untested for the heteropolymeric protein condensates. The success of the computational microrheology technique relies on carefully choosing the parameters for the probe particle, namely its size and interactions with the medium of interest, such that they follow continuum mechanics assumptions. This is also a primary concern in experimental microrheology where the nonspecific interactions between probe beads and protein molecules need to be prevented for reliable measurements.^35^ Establishing a physical basis for choosing the attributes of the probe is important because of the unexplored questions regarding the technique’s capability in two following aspects: (1) capturing the sequence-dependent viscoelasticity of condensates formed by heteropolymeric IDPs and (2) quantifying the spatial variations in viscoelasticity found in heterogeneous condensates formed by a pair of heteropolymeric IDPs. To this end, we systematically demonstrate in this work that computational microrheology is a robust technique for studying the time-dependent rheological properties of protein condensates.

## Results

To investigate the sequence-dependent viscoelasticity of IDP condensates using computational microrheology, we employed coarse-grained (CG) model sequences formed by an equal number of negatively charged glutamic acid (E) and positively charged lysine (K) with chain length *N* = 250 residues. Recently, we used similar IDP sequences to establish the sequence-dependent material properties, namely diffusion coefficient *D* and viscosity *η* of condensates formed by charge-rich model and naturally occurring IDPs.^24^ We chose three E−K sequences that varied in the sequence charge patterning, with a high degree of charge segregation quantified using a high value of normalized sequence charge decoration (nSCD ∈ [0,1]) parameter^42-44^ (see **Table S1** for the amino acid sequences). Specifically, we used variants with nSCD values of 0.067, 0.468, and 1.000, respectively. We performed computational passive probe microrheology via MD simulations in a cubic simulation box at the preferred dense phase concentrations of the E−K variants (see Methods). We found that the dense phase concentration *ρ* increased with increasing degree of charge segregation (i.e., increasing nSCD), highlighting stronger effective interactions between the oppositely charged residues (**Fig. S1**). A probe particle of bare mass *m*_bare_, modeled as a rough sphere for ensuring no-slip boundary conditions,^45^ was dispersed in the dense phase of the E−K variants (**Fig. 1a**). All the simulations were carried out via a transferable hydropathy scale (HPS) model^46, 47^ at a constant temperature of *T* = 300 K (see Methods for model and simulation details).

**Fig. 1.**
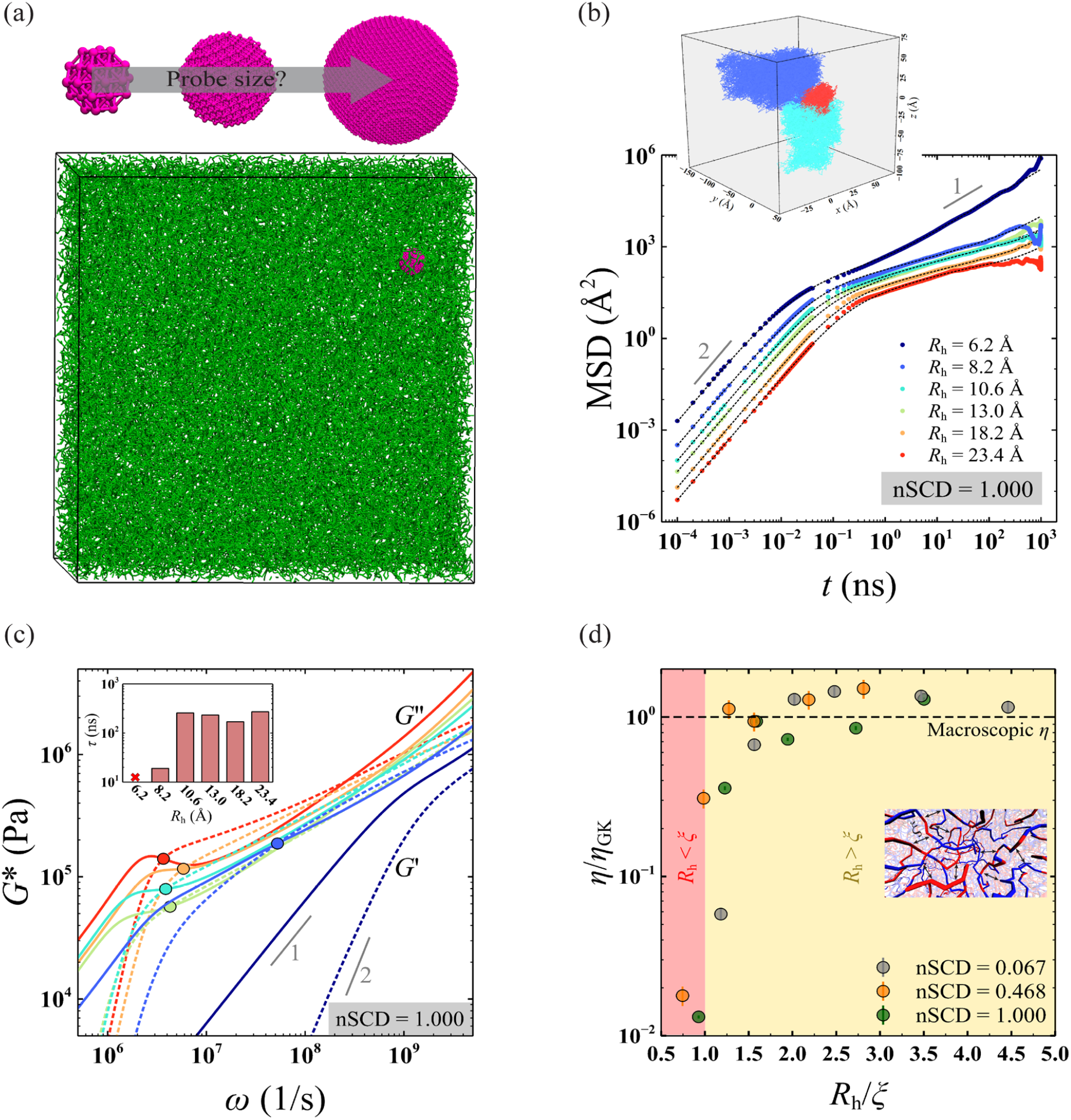
(a) Simulation snapshot of the dense phase of a condensate formed by a select E−K sequence (nSCD = 1) of chain length *N* = 250 with a spherical probe particle (magenta) of hydrodynamic radius *R*_h_ embedded in it. (b) Mean square displacement MSD(*t*) of the center of mass of the probe particle for different *R*_h_ in the dense phase of the E−K sequence with nSCD = 1. The dashed lines are the fits based on the Baumgaertel-Schausberger-Winter-like power law spectrum to the MSD data. The inset shows probe’s displacement in the three-dimensional Cartesian coordinates for select *R*_h_ values. (c) Elastic *G*^′^ (dashed line) and viscous *G*^′′^ (solid line) modulus for the dense phase of the E−K sequence with nSCD = 1 as a function of *R*_h_, with circles corresponding to the crossover frequency. The inset shows the relaxation time *τ*, computed as the inverse of crossover frequency, with varying *R*_h_. (d) Viscosity *η*, normalized by that obtained based on the Green-Kubo (GK) relation *η*_GK_, as a function of normalized probe particle size *R*_h_/*ξ*, where *ξ* is the correlation length, for three different nSCD sequences. The two shaded regions delineate regions where *R*_h_ is smaller or larger than *ξ*. The black dashed line corresponds to *η* = *η*_GK_.

The friction encountered by the probe particle during its Brownian motion can be related to the linear viscoelastic properties (elastic *G*^′^ and viscous *G*^′′^ modulus) of the dense phase of a condensate using the inertial generalized Stokes-Einstein relation (IGSER)^48, 49^

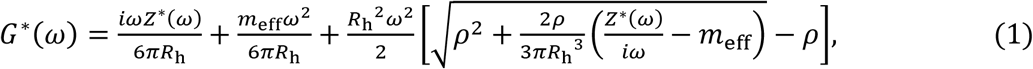

where *G*^*^ = *G*^′^ + *iG*^′′^ is the dynamic modulus of the medium, *Z*^*^ is the frequency-dependent friction experienced by the probe through its interactions with the medium, *R*_h_ is the hydrodynamic radius of the probe, and *m*_eff_ is the effective mass of the probe particle. As per continuum mechanics, *m*_eff_= *m*_bare_+ *m*_add_, where 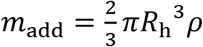 is the added mass from the medium.^50^ For IGSER to accurately capture the viscoelastic properties of IDP condensates, two important characteristics of the probe emerge, namely its size (i.e., *R*_h_) relative to the relevant length scale of the IDP dense phase and its interactions with the IDP chains. This prompted us to do a systematic assessment of the attributes of the probe particle for the successful implementation of computational microrheology for protein condensates.

### Probe particle size is a critical determinant for the computational microrheology of IDP condensates

Given that the length of our model E−K sequences is representative of naturally occurring IDPs, we first asked whether there is an appropriate size of the probe particle, which can be rationalized in terms of the relevant length scale such as the mesh size within IDP condensates.^51^ To investigate this aspect, we varied the probe sizes, ranging from bare radius *R*_b_ = 2.5 Å (similar to the radius of the residue beads in the HPS model) to *R*_b_ = 20 Å (similar to the single-chain radius of gyration *R*_*g*_ of the investigated E−K variants). For our continuum mechanics analysis, we used the hydrodynamic radius *R*_h_ of the probe particle, defined as the location of the first peak in the radial distribution function (RDF) between the probe and the protein residues within the dense phase of the condensates (**Fig. S2**). This choice of definition for *R*_h_ was previously shown to recover the Stokes frictional force and torque for the rough spherical particle moving through a complex fluid as well as yield accurate *G*^′^ and *G*^′′^ values of homopolymers with increasing *N*.^38, 39, 45^ For all sizes of the probe, its interaction with the protein residues was modeled via a modified Lennard-Jones potential with interaction strength *ε*_HPS;_ = 0.2 kcal/mol, the strongest possible van der Waals interaction between a pair of residue beads in our HPS model.

In our passive rheology simulations, we tracked the Brownian displacement of the center of mass of the probe of different sizes in the dense phase of the E−K sequences (insets of **Figs. 1b, S3a, and S4a**). Using the displacement data, we then computed the probe’s mean square displacement

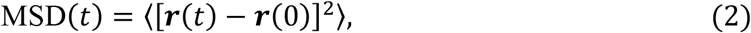

where ***r*** is its position at time *t* (**Figs. 1b, S3a, and S4a**). We found that probes of all sizes *R*_h_ exhibited a ballistic motion (MSD ∝ *t*^2^) at short times, but only the smallest probe (*R*_h_ = 6.2 Å) showed a diffusive motion (MSD ∝ *t*) at long times. We observed a sub-diffusive behavior at intermediate times, which became increasingly prominent until long times with increasing *R*_h_, indicating that larger probes move slower in the condensate within the simulation duration of the E−K sequences investigated. Further, MSD at intermediate to long times drastically decreased for smaller *R*_h_, followed by a gradual decrease for larger *R*_h_, highlighting high friction exerted on large probe particles. In fact, the friction *Z*^*^ that appears in IGSER can be related to the MSD of the probe as

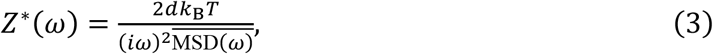

where *d* is the number of dimensions in which the particle is tracked, *k*_B_ is the Boltzmann constant and 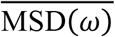 is the one-sided Fourier-transformed MSD. For obtaining MSD in the frequency domain, we used a previously developed analytical fitting procedure by expressing MSD in terms of a Baumgaertel-Schausberger-Winter-like power law spectrum (see Methods; **Figs. 1b, S3a, and S4a**).^39, 52^ This method is usually superior to the use of an approximate Fourier transform expression used for the polynomial fit to MSD(*t*) in experimental microrheology,^37^ which often leads to poor estimates when the slope of MSD varies rapidly.

Next, we used IGSER (Eqs. 1 and 3) to obtain the elastic *G*^′^ and viscous *G*^′′^ modulus of the dense phase of E−K sequences (**Figs. 1c, S3b, and S4b**). Consistent with the trends in the probe motion, we found that *G*^′^ and *G*^′′^ were significantly lower as well as the terminal regime (i.e., *G*^′^ ∝ *ω*^2^ and *G*^′′^ ∝ *ω*^1^) was observed at high frequencies for smaller probe particles (i.e., *R*_h_ ≤ 8.2 Å) as compared to the larger ones for all E−K sequences. For *R*_h_ > 8.2 Å, the onset of terminal viscous characteristics occurred at similar frequency values, as indicated by similar crossover frequencies below which *G*^′′^ > *G*^′^. This was also evident by further looking at the similar relaxation times *τ*, computed as the inverse of crossover frequency, for larger probe particles in the dense phase of E−K variants (insets of **Figs. 1c, S3b, and S4b**). Our findings indicate that for probe sizes beyond a certain threshold value (i.e., *R*_h_ > 8.2 Å), the probe motion yields a viscoelastic spectrum of the dense phase of E−K sequences that exhibit similar characteristics (i.e., similar *τ*).

Given that the motion of the probe is intricately tied to the dense environment of the IDP condensates, we next investigated whether the similar viscoelastic modulus yielded based on the probe motion of certain sizes is because of them being larger than the mesh size^53^ within the dense phase of E−K sequences. Note that IGSER requires the probe particle to be large enough to see the medium as a continuum. Following previous work,^51^ we estimated the correlation length *ξ* from the overlap concentration *ρ*^*^, which is related to the polymer mesh size in a dense phase system. We found the values of *ξ* to be in the range of 5.24 Å to 8.32 Å for the E−K sequences. We also computed the viscosity of the dense phase of E−K sequences from the microrheology simulations as *η* = *G*^′′^/*ω* in the terminal region as well as from the GK relation that yields macroscopic viscosity *η*_GK_ (see Methods). When we normalized *η* by *η*_GK_, we found that probes larger than the correlation length (i.e., *R*_h_/*ξ* ≳ 1.5) yielded macroscopic viscosity (**Fig. 1d**). This finding highlights that probes larger than *ξ* feel the ‘true’ macroscopic friction present within the dense phase. However, given that very large probes sample the dense phase of the condensates much slower (insets of **Figs. 1b, S3a, and S4a**), we concluded that the smallest probe size that satisfied the criteria *R*_h_/*ξ* ≈ 1.5 (i.e., *R*_h_ = 10.6 Å) would be a computationally efficient choice to obtain reliable viscoelastic measurements of any IDP condensates investigated using the HPS model and other analogous CG models.^54-57^

### Confluence of probe particle size and its interaction strength with IDPs correctly captures condensate rheology

The friction experienced by the probe particle depends on its interaction with the IDP chains constituting the condensates. Having established a suitable probe size for computational microrheology, we next investigated whether the strength of probe-protein interactions had a significant effect on the viscoelastic modulus of the dense phase of IDP condensates. An important requirement imposed by IGSER is the need for a no-slip boundary condition at the probe particle surface.^39, 40^ To identify the interaction strengths *ε* that would satisfy the condition, we varied it in the range of *ε*/*ε*_HPS;_ = 0 (purely repulsive interactions) to *ε*/*ε*_HPS;_ = 4 for a probe particle size of *R*_h_ = 10.6 Å. We then computed the velocity *ν*_*x*_ profile of the protein residues around the probe particle that was moving at a pre-defined translational velocity *ν*_*x*,probe_ (**Fig. 2a**). We found that the velocity of the protein residues adjacent to the probe surface became increasingly similar to that of *ν*_*x*,probe_ with increasing *ε*/*ε*_HPS;_, indicating that sufficient attractive interactions prevent a slip at its surface. In line with this observation, we observed that only probe particles with *ε*/*ε*_HPS;_ < 0.75 exhibited very high displacements as well as a diffusive behavior within the course of the simulations of E−K sequences (**Figs. 2b, S5a, and S6a**). To quantify the variations in the probe’s MSD for different *ε*/*ε*_HPS;_ in terms in viscoelasticity, we measured *G*^′^ and *G*^′′^ of the dense phase of E−K variants via IGSER (**Figs. 2c, S5b, and S6b**).

**Fig. 2.**
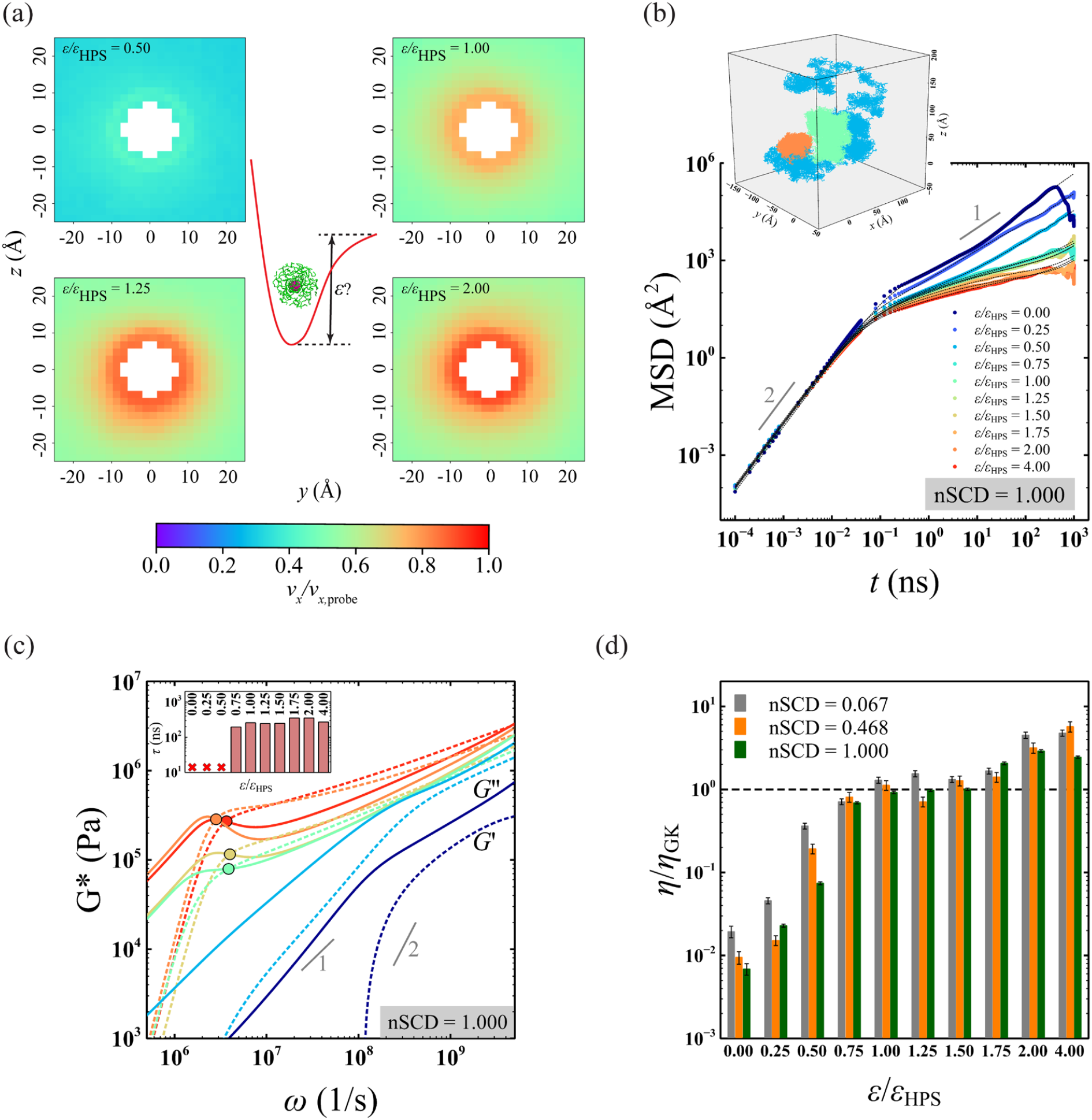
(a) Velocity *ν*_*x*_ profile of the residues of protein chains with nSCD = 1 around the probe particle translating with a velocity *ν*_*x*,probe_ for different probe-protein interaction strengths *ε*, normalized by the strongest possible interaction strength *ε* = 0.20 kcal/mol in the HPS model. (b) Mean square displacement MSD(*t*) of the center of mass of the probe particle for different *ε*/*ε*_HPS;_ in the dense phase of the E−K sequence with nSCD = 1. The dashed lines are the fits based on Baumgaertel-Schausberger-Winter-like power law spectrum to the MSD data. The inset shows probe’s displacement in the three-dimensional Cartesian coordinates for select *ε*/*ε*_HPS;_ values. (c) Elastic *G*^′^ (dashed line) and viscous *G*^′′^ (solid line) modulus for the dense phase of the E−K sequence with nSCD = 1 as a function of *ε*/*ε*_HPS;_, with circles corresponding to the crossover frequency. The inset shows the relaxation time *τ* with varying *ε*/*ε*_HPS;_. (d) Viscosity *η*, normalized by that obtained based on the Green-Kubo (GK) relation *η*_GK_ as a function of *ε*/*ε*_HPS;_ for three different nSCD sequences. The black dashed line corresponds to *η* = *η*_GK_.

For probes with *ε*/*ε*_HPS;_ < 0.75, we found that *G*^′^ and *G*^′′^ were significantly lower and did not show a crossover between them as compared to the higher *ε*/*ε*_HPS;_ values. When sufficient attractive interactions (*ε*/*ε*_HPS;_ ≥ 0.75) exist between the probe and the protein residues, similar relaxation timescales delineating dominant viscous and elastic behaviors were observed irrespective of *ε*/*ε*_HPS;_ for all sequences (**Figs. 2c, S5b, and S6b**). However, for *ε*/*ε*_HPS;_ = 4, the modulus values were much higher as compared to *ε*/*ε*_HPS;_ = 1. This is further highlighted in the normalized viscosity *η*/*η*_GK_ values, which revealed an optimal window for interaction strengths (1 ≤ *ε*/*ε*_HPS;_ ≤ 1.5) that yielded the macroscopic viscosity (**Figs. 2d**). Note that, for *ε*/*ε*_HPS;_ ≳ 1.75, we observed the IDP chains getting strongly adsorbed to the probe particle surface (**Fig. S7**). This is a common concern in experimental microrheology as well, in which, for example, polystyrene beads are often passivated with polyethylene glycol, to ensure negligible chemical interactions with the biomolecules as well as to prevent the beads from getting constrained within the droplet.^35, 58, 59^ From these findings, we concluded that an attractive strength (*ε*/*ε*_HPS;_ = 1) that is low enough, but a value that resides in the optimal window would be ideal for correctly capturing the time-dependent rheology of IDP condensates computationally.

### Computational microrheology reveals the sequence-encoded time-dependent viscoelasticity of heteropolymeric IDP condensates

Through the continuum analysis of the motion of a single probe particle in our computational microrheology simulations, we have identified suitable probe particle size and its interaction strength with the IDP chains that would yield accurate macroscopic dense phase viscosities of the E−K sequences. However, the microrheology experiments are often performed with multiple probe particle beads within the same droplet, and the average motion of all particles is used for quantifying the condensate viscoelasticity.^13, 28^ This approach is efficient as it reduces the uncertainties in the viscoelastic measurements, which may arise from tracking only a single particle. Though the continuum mechanics expressions are for a single particle, which necessitates that no hydrodynamic interactions exist between the probe particles, we next asked the extent to which multiple probes in an IDP dense phase system would influence their motion, which can alter the corresponding viscoelastic measurements. For this purpose, we performed the passive rheology simulation with *n* = 2 to 48 probes in the dense phase of E−K sequences (**Fig. S8**). Surprisingly, we found that the probe’s average MSD was nearly the same at all times with varying *n*, but the statistical noise in the MSD profile has significantly reduced for systems with multiple particles. To demonstrate the ability of passive rheology simulations to capture the changes in viscoelasticity with increasing nSCD, we again obtained the viscoelastic modulus *G*^′^ and *G*^′′^ of the dense phase of E−K sequences based on the average MSD of *n* = 8 probes (**Fig. 3a**). The modulus values obtained from the microrheology simulations using multiple probes are in quantitative agreement with those obtained based on a single probe particle (**Fig. S9**). Further, we also observed a semi-quantitative agreement, yet similar trends in the modulus values obtained from the microrheology simulations and based on the Green-Kubo relation for all E−K sequences, even though the latter method suffers from noise at long times (see Methods; **Figs. S10 and S11**). Our microrheology simulations revealed that the modulus *G*^′^ and *G*^′′^, relaxation time *τ*, and viscosity *η* were higher with increasing nSCD, highlighting that charge segregation slows down dynamics, thereby resulting in longer timescales for displaying the terminal flow behavior (**Fig. 3a**). Finally, we also tracked the probe motion in the dense phase of E−K sequences of *N* = 50, but with the same nSCD values as those used for *N* = 250 (**Fig. S12a**). We found that *G*^′^ and *G*^′′^ did not show a cross-over (i.e., no dominant elastic response in the entire frequency space) for the shorter sequences, indicating negligible entanglement effects between the chains in these systems (**Fig. S12b**). Our findings indicate that condensates formed by longer chains exhibit Maxwell fluid-like behavior over the entire frequency range investigated, which is in agreement with recent experimental^28^ and computational studies.^60, 61^

**Fig. 3.**
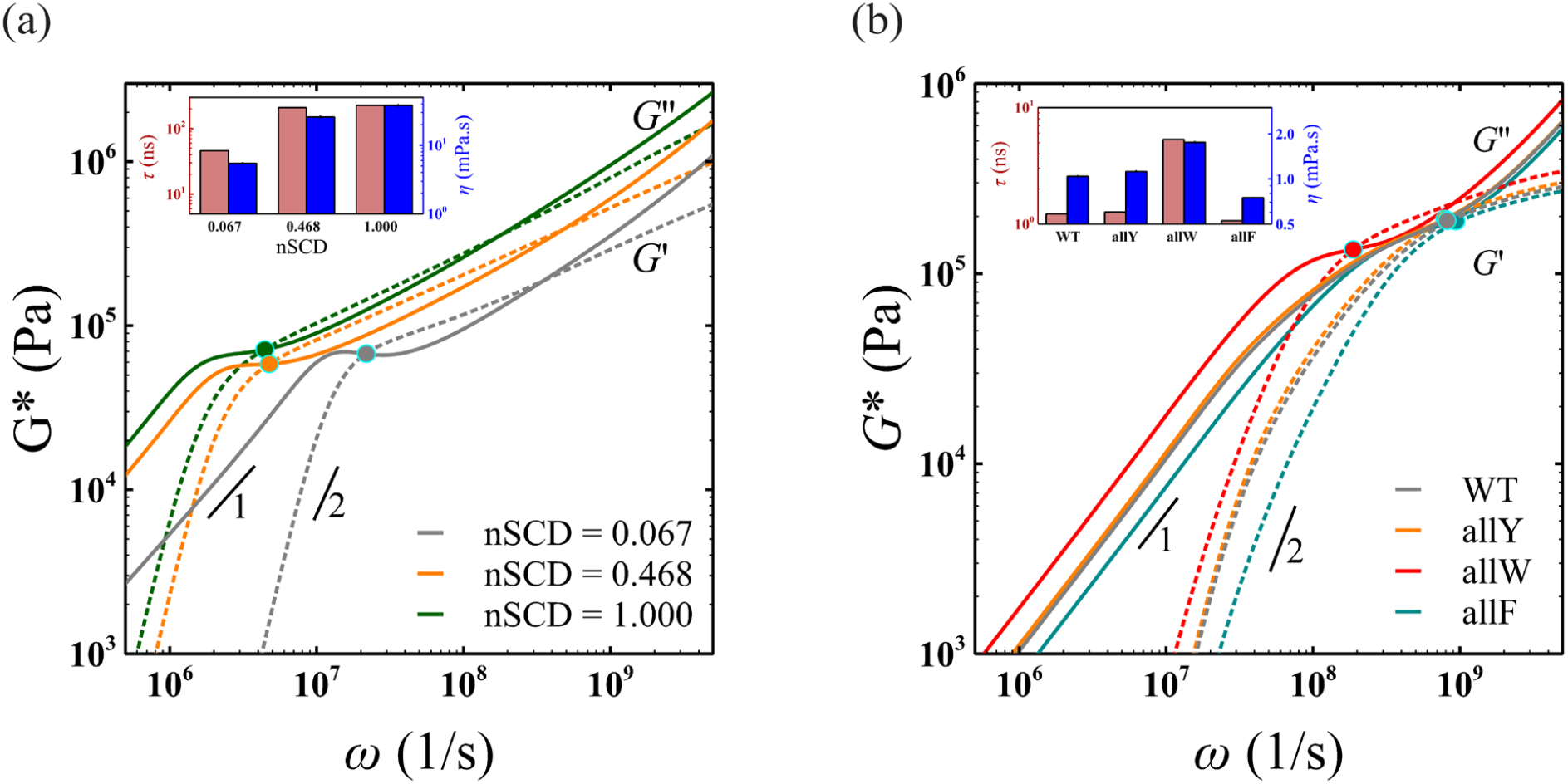
(a) Elastic *G*^′^ (dashed line) and viscous *G*^′′^ (solid line) modulus for three E−K sequences with different nSCD, obtained based on multiple (*n* = 8) probes in each dense phase system. (b) Elastic *G*^′^ (dashed line) and viscous *G*^′′^ (solid line) modulus for A1-LCD WT and its three variants, obtained based on multiple (*n* = 6) probes in each dense phase system. The circles in (a) and (b) correspond to the sequence-specific crossover frequencies. The insets in (a) and (b) shows the changes in relaxation time *τ* and viscosity *η* with changing nSCD and with mutational changes in the A1-LCD WT sequence, respectively.

The dynamics and rheology of IDP condensates can be modulated not only through sequence charge patterning but also via alterations to sequence hydrophobicity. We next investigated the ability of the passive rheology technique coupled with our CG model to capture the sequence alterations affecting the hydrophobic character. To this end, we tracked multiple (*n* = 6) probe particles in a naturally occurring IDP A1-LCD wildtype (WT) sequence and three of its variants, each with *N* = 137, with different aromatic residue (tyrosine Y, tryptophan W, phenylalanine F) identities, namely allY, allW, and allF (see **Table S2** for the amino acid sequences). Again, we ensured that the average probe MSD computed based on *n* = 6 particles was similar to the MSD of a single particle in these systems, except for long times where the statistics for the MSD based on multiple probes were better as compared to a single probe MSD (**Fig. S13**). The values of *G*^′^ and *G*^′′^, *τ*, and *η* obtained using IGSER increased in the following order: allW > allY ≈ WT > allF (**Fig. 3b**). This finding is in agreement with the recent experimental microrheology measurements on the same set of IDP sequences.^62^ Taken together, we concluded that the computational microrheology technique coupled with continuum mechanics and CG models can accurately quantify the sequence-encoded time-dependent viscoelasticity of IDP condensates.

### Computational microrheology unmasks the spatial variations in the viscoelasticity of heterogeneous IDP condensates

Intracellular biomolecular condensates are multi-component in nature, often exhibiting complex molecular architectures with spatial heterogeneities.^63^ We next asked whether computational microrheology can be used to quantify the spatial variations in the viscoelasticity of heterogeneous condensates. It is known that IDPs with large differences in nSCD values demix in such a way that a highly charge-segregated sequence forms the condensate core and the well-mixed sequence forms a shell around the core.^64^ Specifically, we formed such a condensate using a pair of E−K sequences with nSCD = 0.067 and nSCD = 1.000, respectively (**Fig. 4a**). Also, the maximum densities of the two E−K sequences in the heterogeneous mixture were the same as that we observed in the bulk condensates simulated in a cubic geometry (**Fig. 4a**). To characterize the local viscoelasticity within this heterogeneous condensate, we spatially restrained *n* = 3 probes along the *z* direction encompassing distinct local environments: one among nSCD = 1.000 chains, one among nSCD = 0.067 chains, and one at the interface of the two nSCD sequences (see Methods). We found that the two-dimensional probe particle displacements were different in each of the three regions at intermediate to long times, with the probe’s among nSCD = 0.067 chains and nSCD = 1.000 chains exhibiting the highest and lowest mobilities, respectively (**Fig. 4b**). This was evident in the elastic *G*^′^ and viscous *G*^′′^ modulus obtained by using the two-dimensional probe particle displacement information in IGSER (**Fig. 4c**). Specifically, the values of modulus and the corresponding relaxation times *τ* were lower for the shell region around the condensate core as compared to the core itself, while the values at the interface between the core and the shell regions displayed intermediate viscoelasticity. Interestingly, we also found that viscosity *η* values at the condensate core and shell regions, obtained based on *G*^′′^ in the terminal region, were quantitatively similar to that obtained from the bulk condensates (**Fig. 4d**). These observations further highlight the potential of computational microrheology for revealing the location-dependent viscoelasticity of complex heterogeneous condensates, which can gain insights about the small molecules that partition into such environments^27^ as well as help define the role of condensate interfaces that have been found to exhibit distinct conformational characteristics as compared to the core of the condensates.^65, 66^

**Fig. 4.**
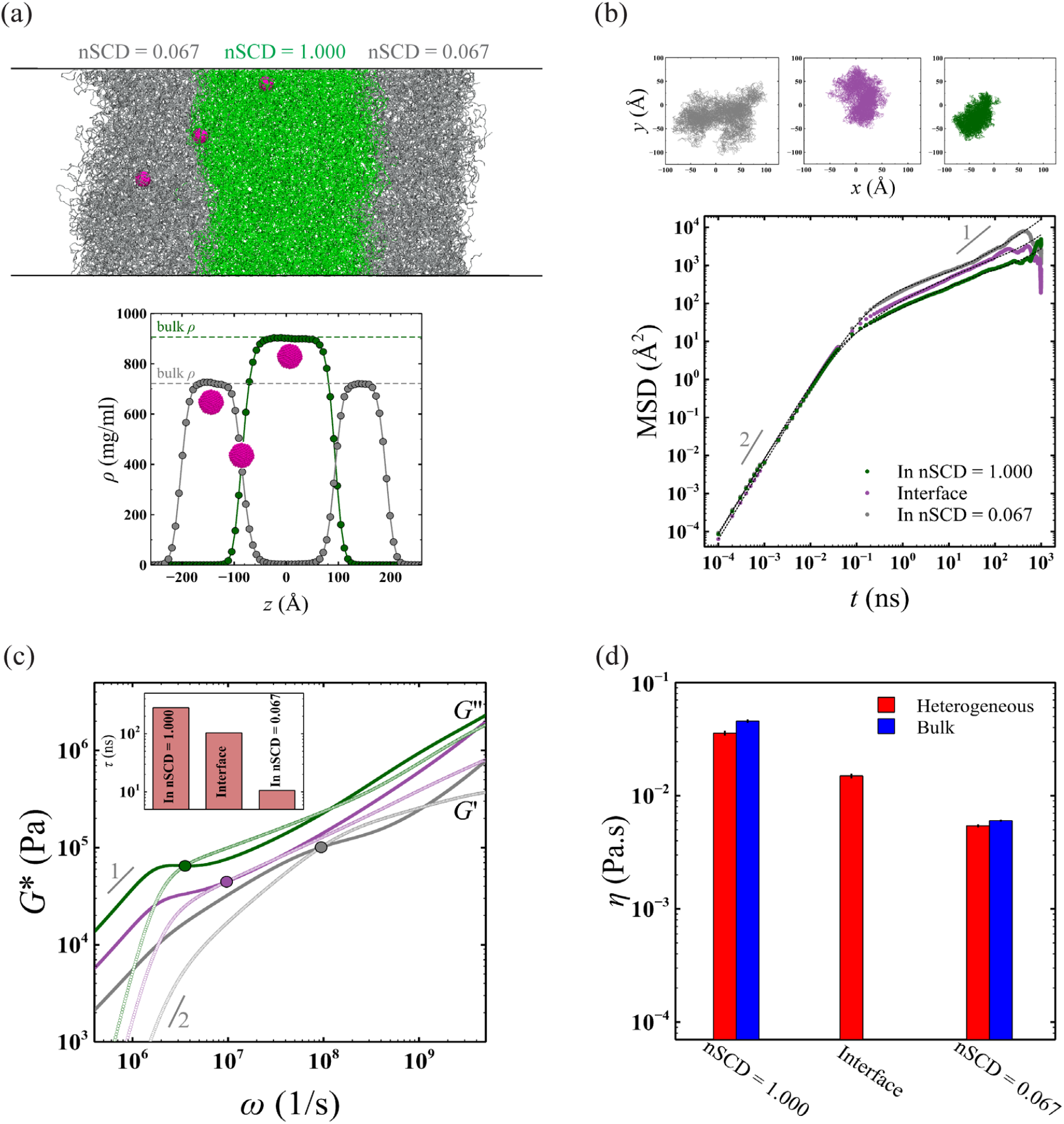
(a) Simulation snapshot of a heterogeneous condensate formed by two different E−K sequences with nSCD = 0.067 and nSCD = 1.000. Three probe particles are spatially restrained in three regions of the heterogeneous condensate: one among nSCD = 1.000 chains, one among nSCD = 0.067 chains, and one at the interface of the two nSCD sequences. Also shown are the density profiles of the heterogeneous condensate. (b) Mean square displacement MSD(*t*) of the probe particle in three different regions of the heterogeneous condensate. The dashed lines are the fits based on Baumgaertel-Schausberger-Winter-like power law spectrum to the MSD data. The inset shows probe’s displacement in the two-dimensional Cartesian coordinates in the chosen regions within the condensate. (c) Elastic *G*^′^ (open symbols) and viscous *G*^′′^ (closed symbols) modulus sampled from three different regions of the heterogeneous condensate, with circles corresponding to the location-specific crossover frequency. The inset shows the relaxation time *τ* in the chosen regions within the condensate. (d) Viscosity *η* in three regions of the heterogeneous condensate compared with those obtained from the bulk simulations of the E−K sequences.

## Conclusions

Knowledge of the molecular interactions that govern the viscoelastic transitions would allow for establishing how protein sequence dictates the spatiotemporal evolution of condensates. Using the HPS model, we demonstrate that a computational passive probe microrheology technique, in which the probe particle motion is analyzed via continuum mechanics, can accurately quantify the sequence-dependent viscoelasticity of IDP condensates. We do so by first rationalizing our choice for the two important attributes of the probe particle, namely its size and interaction strength with the IDPs. We found that a probe with a hydrodynamic radius *R*_h_ being slightly greater than the correlation length *ξ* within the condensates (*R*_h_ ≈ 1.5*ξ*) and its interactions with the IDPs being optimally strong that it prevents slip (*ε*/*ε*_HPS;_ = 1) is sufficient to accurately quantify the viscosity and viscoelasticity of IDP condensates. By tracking the motion of the probe particle and converting its displacement information into viscoelastic modulus using IGSER, we found that the measurements from microrheology simulations are in quantitative agreement with those obtained based on the Green-Kubo relation. Further, the elastic and loss modulus, relaxation time, and terminal viscosity increased with increasing degree of charge segregation, exhibiting Maxwell fluidic nature for the E−K sequences with identical sequence composition. This observation highlighted that computational microrheology captures the time-dependent rheological transitions with sequence alterations that result in pronounced electrostatic interactions between the oppositely charged residues. Further, we have shown that the microrheology simulations can capture changes in viscoelasticity with changes in the composition of aromatic residues in the naturally occurring A1-LCD WT sequence through mutations. Taken together, we conclude that computational microrheology is a robust technique for characterizing the sequence-encoded time-dependent viscoelasticity of heteropolymeric protein condensates.

Microrheology experiments are often performed with multiple probe particles within the same droplet and the average displacement of the particle is used to characterize condensate viscoelasticity. We also showed that microrheology simulations performed with multiple probe particles yield accurate viscoelastic modulus values of the dense phase of IDP sequences investigated in this work. This suggests that a single simulation with multiple probes can lead to improved statistical accuracy of the probe’s displacement at long times. Consequently, condensate viscoelasticity can be sampled based on a single simulation trajectory as opposed to other conventional techniques such as the MD simulations along with the Green-Kubo (GK) relation and the NEMD simulations. In the method involving GK relation, the values of shear stress relaxation modulus, used for obtaining the viscoelastic modulus, are typically prone to noise at long times because of which very long MD simulations are required.^22^ In the NEMD method, viscoelastic modulus can only be obtained by applying an oscillatory shear strain on the system at a specific frequency and is prohibitive of accessing low frequency (long time) viscoelasticity, rendering it computationally expensive.^26, 38^ Further, the use of multiple probe particles can allow for the accurate characterization of viscoelasticity in biomolecular condensates with inherent heterogeneities due to physical aging.^62, 67^ Finally, we spatially restrained multiple probe particles at distinct locations within a heterogeneous IDP condensate formed by a pair of E−K sequences with vastly different charge patterning for quantifying local viscoelasticity. By tracking each of the particle’s displacements in unrestrained directions, we demonstrated the technique’s ability to accurately quantify the viscoelasticity that depends on the local environment within such condensates. This finding highlights that passive rheology can be used for accurately sampling the spatial variations in viscoelasticity that are prevalent in intracellular condensates. Knowledge of the spatial dependence of viscoelasticity can provide important insights for designing drug-delivery nanoparticles with targeted partitioning and transport within condensates.^68^ We believe that the computational microrheology technique can be an easy-to-use technique for establishing the sequence determinants of condensate viscoelasticity for a wide range of protein condensates.

## Methods

### Hydropathy scale (HPS) model

We used our transferable CG model based on the hydropathy scale to computationally investigate the IDP sequences.^46, 47^ This CG framework has been widely used in deciphering the sequence-dependent conformations and phase separation of a wide range of IDPs.^44, 54, 69-74^ In this framework, we represent the IDPs as fully flexible polymeric chains with a single bead per residue representation. Interactions between bonded residues occurred via the harmonic potential

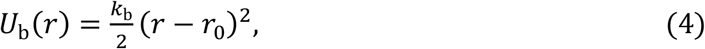

with distance *r* between residues, spring constant, and equilibrium bond length set to *k*_b_ = 20 kcal / (mol Å^2^) and *r*_0_ = 3.8 Å, respectively. The van der Waals interactions between nonbonded residues *i* and *j* were modeled using the modified Lennard-Jones (LJ) potential^75, 76^ based on the average hydropathy *λ* = *λ*_*i*_ + *λ*_*j*_/2

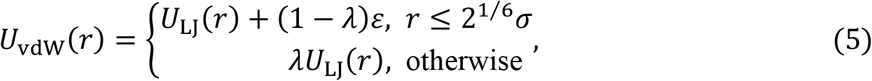

where *U*_LJ_ is the standard LJ potential

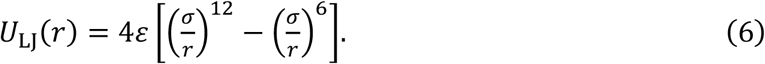

The parameter of LJ potential are the average diameter *σ* = (*σ*_*i*_ + *σ*)/2, and the interaction strength *ε* = *ε* _HPS;_ = 0.2 kcal/mol. We used *λ* values based on the Kapcha-Rossky scale^77^ for the E−K sequence variants and those based on the Urry scale^78^ for the A1-LCD sequence variants. We used the Urry scale for A1-LCD as it is known to capture the changes in the phase behavior of natural proteins upon mutations of arginine to lysine and tyrosine to phenylalanine.^47^ The values of *U*_vdW_ and its forces were truncated to zero at a distance of 4 *σ*. Finally, the nonbonded charged residues interacted through a Coulombic potential with Debye-Hückel electrostatic screening^79^

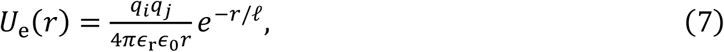

with vacuum permittivity *∈*_0_, relative permittivity *∈*_r_ = 80, and Debye screening length *ℓ* = 10 Å. The choices for *∈*_r_ and *ℓ* were made to represent an aqueous solution with a physiological salt concentration of ∼100 mM. The values of *U*_e_ and its forces were truncated to zero at a distance of 35 Å.

### Model of the probe particle and its interactions with IDPs

We modeled the probe particle by carving out a spherical region from a face-centered cubic (FCC) crystal lattice structure of the LJ beads (*σ* = 1.5 Å and mass *m* = 100 g/mol), with a lattice spacing (i.e. distance between corner atoms) of 2.12 Å. This value was chosen because it ensures that the corner and the face atoms in the FCC lattice are at a distance of 1.5 Å (i.e., just touching each other). The spherical shape of the probe particle was maintained by connecting the neighboring LJ particles, constituting the corner and the face atoms, using stiff harmonic bonds with a spring constant *k*_b_ = 250 kcal/mol Å^2^. We ensured that *R*_g_ of the probe particle was nearly identical to the expected value for a spherical particle 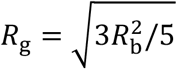 as well as its relative shape anisotropy *κ*^2^ was nearly zero (*κ*^2^ = 0 for a sphere) during the simulations (**Fig. S14**).^53^

The probe particle beads interacted with all protein residues in an IDP chain via the modified LJ potential *U*_vdW_ (Eqs. 5 and 6), in which the values of *λ* were varied between the values of 0 and 4 to control the interaction strength *ε*. Specifically, the values of *ε* were in the range of 0 kcal/mol to 0.8 kcal/mol. Again, we truncated *U*_vdW_ and its forces were to zero at a distance of 4 *σ*. We note that the interactions between the beads constituting the probe particle were modeled using a purely repulsive potential, which corresponds to *λ* = 0 in Eq. 5.

### Microrheology simulation details

We simulated the IDP sequences in a cubic simulation box at a constant pressure of *P* = 0 atm for a duration of 0.2 *μs*. At the end of this equilibration run, the IDPs reached their preferred sequence-dependent dense phase concentration *ρ*. We then performed the Langevin dynamics (LD) simulations in the canonical ensemble for a total duration of 1 *μs*. For these simulations, a damping factor of *t*_damp_ = 1 ns was used to set the friction coefficient of a residue in the chain as well as a bead constituting the probe particle to *f* = *m*/*t*_damp_.

For characterizing the spatial dependence of viscoelasticity, we performed LD simulations of a heterogeneous condensate formed by a pair of E−K sequences in a slab geometry (225 Å × 225 Å × 1687.5 Å) for a duration of 1 *μs*. The friction coefficient for the IDP residues and the probe particle beads were the same as those used in the bulk dense phase simulations. Three probe particles were restrained via a harmonic potential with *k*_b_ = 20 kcal/mol Å^2^ at specific locations along the *z* direction of the heterogeneous condensate via the restrain functionality (i.e., restrain.plane) within azplugins.^80^

For comparison with the microrheology simulations, we performed equilibrium MD simulations of the E−K sequence variants in the absence of a probe particle to compute their viscosity and viscoelasticity using the Green-Kubo relation

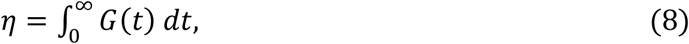

where *G*(*t*) is the shear stress relaxation modulus. We measured *G*(*t*) (**Fig. S15**) based on the autocorrelation of the pressure tensor components *P*_*ab*_^22, 81^

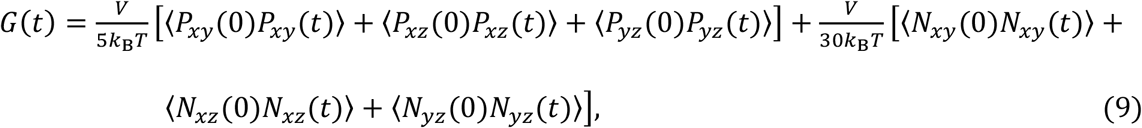

where *V* is the volume of the simulation box and *N*_*ab*_ = *P*_*ab*_ − *P*_*ab*_ is the normal stress difference. For computing viscosity *η*, we followed the approach of Tejedor et al.^22^ by fitting the smooth *G*(*t*) profile at long times to a series of Maxwell modes (*G*_*i*_exp (−*t*/*τ*_*i*_) with *i* = 1….4) equidistant in logarithmic time.^53^ We obtained *η* by summing up the values from numerical integration at short times and analytical integration based on the fits to Maxwell modes at long times. For computing elastic *G*^′^ and loss *G*^′′^ modulus, we Fourier transformed *G*(*t*) using the RepTate software.^82^

All the simulations were performed with periodic boundary conditions applied to all three Cartesian directions. The simulations were performed with a timestep of 10 fs using HOOMD-blue (version 2.9.3)^83^ with features extended using azplugins (version 0.10.1).^80^

### Continuum analysis of the probe motion

In microrheology, the viscoelastic modulus (elastic *G*^′^ and loss *G*^′′^ modulus) is estimated from the probe particle’s motion, which is considered to obey the generalized Langevin equation (GLE)

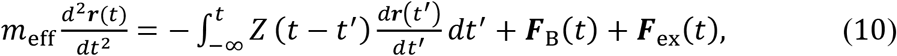

where *Z*(*t*) is the time-dependent friction, ***F***_B_ is the Brownian force on the probe particle, and ***F***_ex_ is the external force on the probe particle, which is zero in the case of our passive rheology simulations. Because of the low time scales inherent in the MD simulations, inertia plays an important role in accurately quantifying the modulus values of protein condensates. When the inertial terms are included, the GLE in the frequency domain takes the form^38, 39, 48^

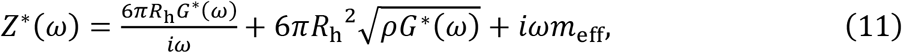

where the terms on the right side correspond to the generalized Stokes drag, the Basset force arising from the medium inertia, and the effective probe particle inertial force, respectively. In experimental microrheology, the generalized Stokes drag alone is sufficient to obtain the viscoelastic modulus of complex fluid systems. On rearranging Eq. 11, we obtained IGSER as given in Eq. 1.

### Analytical expression describing the probe’s displacement data

We followed the approach originally proposed by Karim et al.^38, 39^, which we discuss here for the sake of completeness. The probe’s mean square displacement can be described through the Baumgaertel-Schausberger-Winter-like power law spectrum

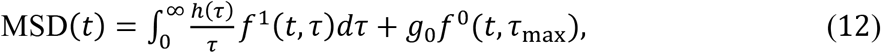

with *f*^1^, *f*^0^, *τ* being the first function for capturing the short-time ballistic behavior (MSD = *C*t^2^, where *C* is the ballistic coefficient), the second function for capturing the long-time diffusive behavior (MSD = 6*Dt*, where *D* is the diffusion coefficient), and characteristic time representing the changes in MSD, respectively. Further, the spectrum *h*(*τ*) can be written as

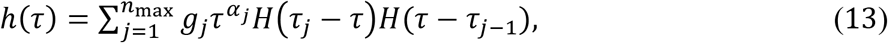

where *n*_max_ is the number of *j* modes, with each mode having a relaxation time *τ*_*j*_ (*τ*_*j*_ > *τ*_*j*−1_) and exponent *α*_*j*_. *H*(*τ*) is the Heaviside step function. The function in the first term is

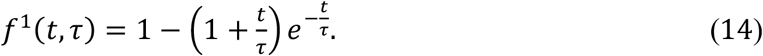

The function in the second term is

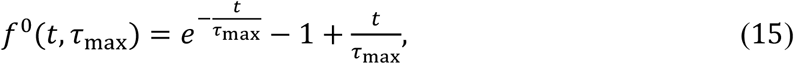

which is weighted by the constant 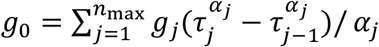, ensuring that the first term in Eq. (12) equals the second term when *t* = *τ*_max_. The other weighted terms include 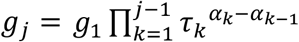 for 2 ≤ *j* ≤ *n*_max_. The value of *g*_1_ can be obtained from 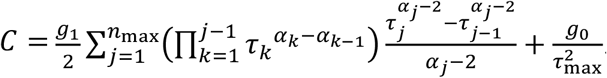. Given that *τ*_max_ is usually large, the second term in the expression for *C* is negligible, and once the slope of the ballistic regime is known, *g*_1_ and other weighted terms can be readily computed. The integration of Eq. (12) with respect to *τ* then gives rise to

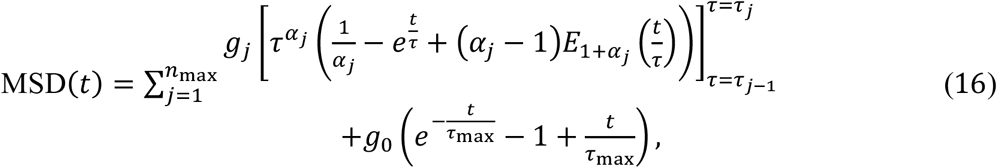

where 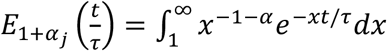 is the exponential integral function. The Fourier transform of Eq. (16) gives the real and imaginary parts of the MSD in the frequency domain as

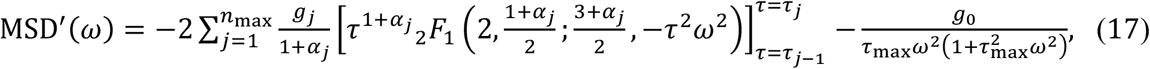

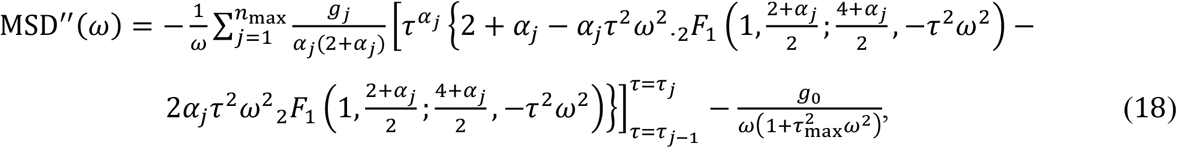

where ._2_*F*_1_ is the hypergeometric function and 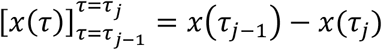 for an arbitrary function *x*(*τ*). The parameters {*α*_1_, …, *α*_max,_*τ*_0_, *τ*_1_, …, *τ*_max_} are obtained based on the fit to the probe’s MSD data. We fixed *τ*_*j*_ values and numerically sought *α*_j_ values that minimize *χ*^2^ of MSD, which were then used to obtain the MSD values in the frequency domain (Eqs. 17 and 18).

## Acknowledgments

This material is based on the work supported by the National Institute of General Medical Science of the National Institutes of Health under grants R01GM136917 and R35GM153388, and the Welch Foundation under grant A-2113-20220331. We thank Prof. Michael P. Howard (Auburn University) for bringing to our attention about the features available within azplugins to restrain the particles at specific locations in our simulations. We also thank Prof. Benjamin Schuster (Rutgers University) and Dr. Avijeet Kulshrestha (Texas A&M University) for their helpful comments on the manuscript. The authors acknowledge the Texas A&M High Performance Research Computing (HPRC) for providing computational resources that have contributed to the results reported in this research article.

