## Supplementary Information for "Sequence-encoded Spatiotemporal Dependence of Viscoelasticity of Protein Condensates Using Computational Microrheology"

**Table S1.** Amino acid codes of E−K sequence variants of chain length $N = 250$ that were used in this study.

| E−K | Sequence |
|---|---|
| nSCD = 0.067 | EEKEEEEKEKEEEEKEEKKEKKKEEKEKEEKKEKEKEKEKEEEEEKKEEEEEEEEE<br>KEKEEEEKKKKKKEEEEEKKKEEKKKEKKEEEKEEEKEKKEEKEEEEEEEEKEEEE<br>EKKEEKEKEEKKKEKEEKEKEEKKEKKKEKKKKEEEKKKEKEKKKEKKKEEKEEKE<br>KEKKKKEKEKEKKEKEKKEKEKKKKEKKKKEEKEKKEEEKKKEKEKKKKKKEKEKK<br>KKKEEKKKKKKEEEKKEEKKKEKKEE |
| nSCD = 0.468 | EEEEEEEEEEEEEEKEEEEEEEEEEEKEEEEEEEEEEEEEKEEEEEEEEKEEEEEE<br>EKEEEEEEKEEKKKEEKEKEKEEEEKEKEEEEEKEEEEKKEEEEEEEEKEEEEEEE<br>EEEEEKKEKKEKKKEEKEKKEKKEKKKKKEKKKKEKKKKKKEKKKKKKEKEKEKEK<br>KEKKKKKKKKKEEKKKKEKKKKKKKKKKKEEKKKKKKEKKKKKKKKKEKKKKKKKK<br>KKKKKKKEKEKKKKKKKKEEKKKKKK |
| nSCD = 1.000 | EEEEEEEEEEEEEEEEEEEEEEEEEEEEEEEEEEEEEEEEEEEEEEEEEEEEEEEE<br>EEEEEEEEEEEEEEEEEEEEEEEEEEEEEEEEEEEEEEEEEEEEEEEEEEEEEEEE<br>EEEEEEEEEEEEEKKKKKKKKKKKKKKKKKKKKKKKKKKKKKKKKKKKKKKKKKKK<br>KKKKKKKKKKKKKKKKKKKKKKKKKKKKKKKKKKKKKKKKKKKKKKKKKKKKKKKK<br>KKKKKKKKKKKKKKKKKKKKKKKKKK |

**Table S2.** Amino acid codes of A1-LCD wildtype (WT) and its sequence variants of chain length $N = 137$ that were used in this study.

| A1-LCD | Sequence |
|---|---|
| **WT** | GSMASASSSQRGRSGSGNFGGGRGGGFGGNDNFGRGGNFSGRGGFGGSRGGGGYGG SGDGYNGFGNDGSNFGGGGSYNDFGNYNNQSSNFGPMKGGNFGGRSSGPYGGGGQY FAKPRNQGGYGGSSSSSSYGSGRRF |
| **allY** | GSMASASSSQRGRSGSGNYGGGRGGGYGGNDNYGRGGNYSGRGGYGGSRGGGGYGG SGDGYNGYGNDGSNYGGGGSYNDYGNYNNQSSNYGPMKGGNYGGRSSGPYGGGGQY YAKPRNQGGYGGSSSSSSYGSGRRY |
| **allW** | GSMASASSSQRGRSGSGNWGGGRGGGWGGNDNWGRGGNWSGRGGWGGSRGGGGWGG SGDGWNGWGNDGSNWGGGGSWNDWGNWNNQSSNWGPMKGGNWGGRSSGPWGGGGQW WAKPRNQGGWGGSSSSSSWGSGRRW |
| **allF** | GSMASASSSQRGRSGSGNFGGGRGGGFGGNDNFGRGGNFSGRGGFGGSRGGGGFGG SGDGFNGFGNDGSNFGGGGSFNDFGNFNNQSSNFGPMKGGNFGGRSSGPFGGGGQF FAKPRNQGGFGGSSSSSSFGSGRRF |

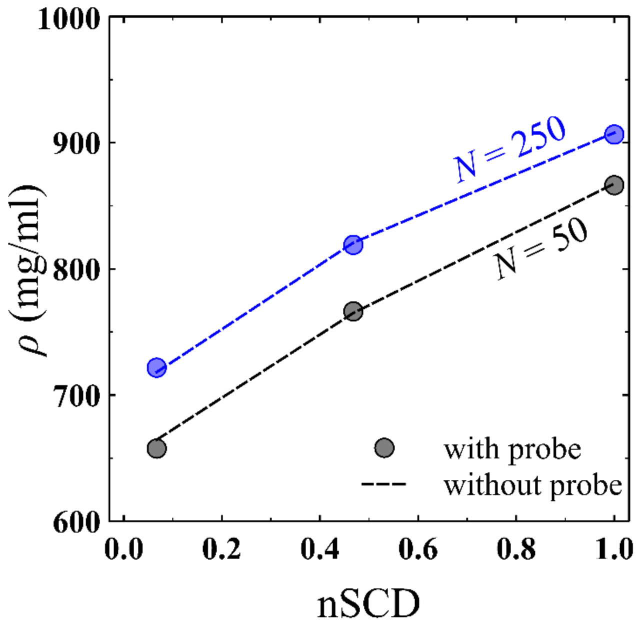


**Fig. S1.** Dense phase concentration $\rho$ attained by the E−K sequence variants of chain lengths $N = 50$ and $N = 250$ with and without the presence of a probe particle in the system.

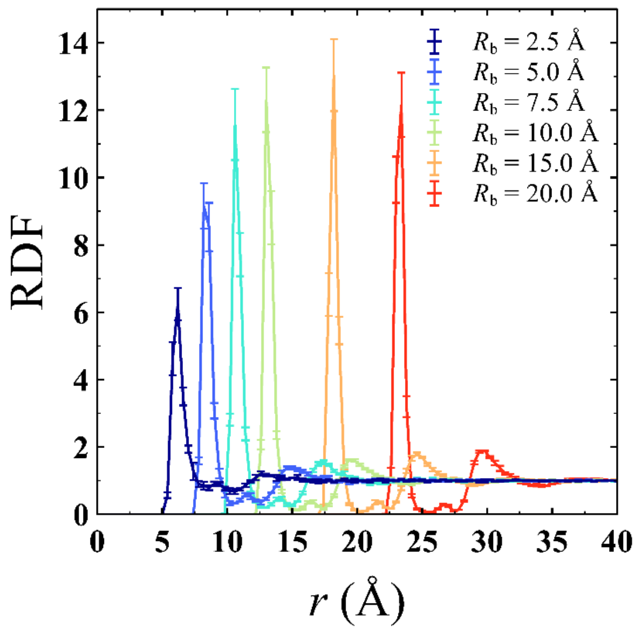


**Fig. S2.** Radial distribution function RDF between the center of mass of the probe and the residues of the protein chains as a function of probe's bare radius $R_b$ for the E−K sequence variants of chain length $N = 250$.

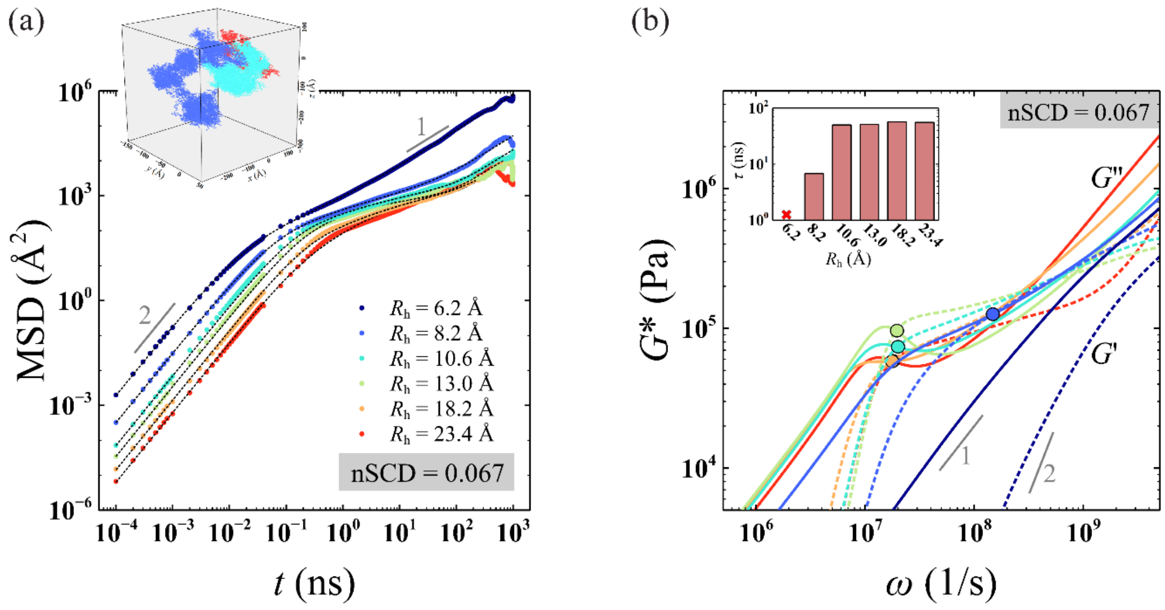


**Fig. S3.** (a) Mean square displacement MSD$(t)$ of the probe particle of different hydrodynamic radius $R_{\mathrm{h}}$ in the dense phase of the E−K sequence with nSCD = 0.067. The dashed lines are the fits based on the Baumgaertel-Schausberger-Winter-like power law spectrum to the MSD data. The inset shows probe particle's displacement in the three-dimensional Cartesian coordinates for select $R_{\mathrm{h}}$ values. (b) Elastic $G'$ (dashed line) and viscous $G''$ (solid line) modulus for the dense phase of the E−K sequence with nSCD = 0.067 as a function of $R_{\mathrm{h}}$, with circles corresponding to the crossover frequency. The inset shows the relaxation time $\tau$ with varying $R_{\mathrm{h}}$.

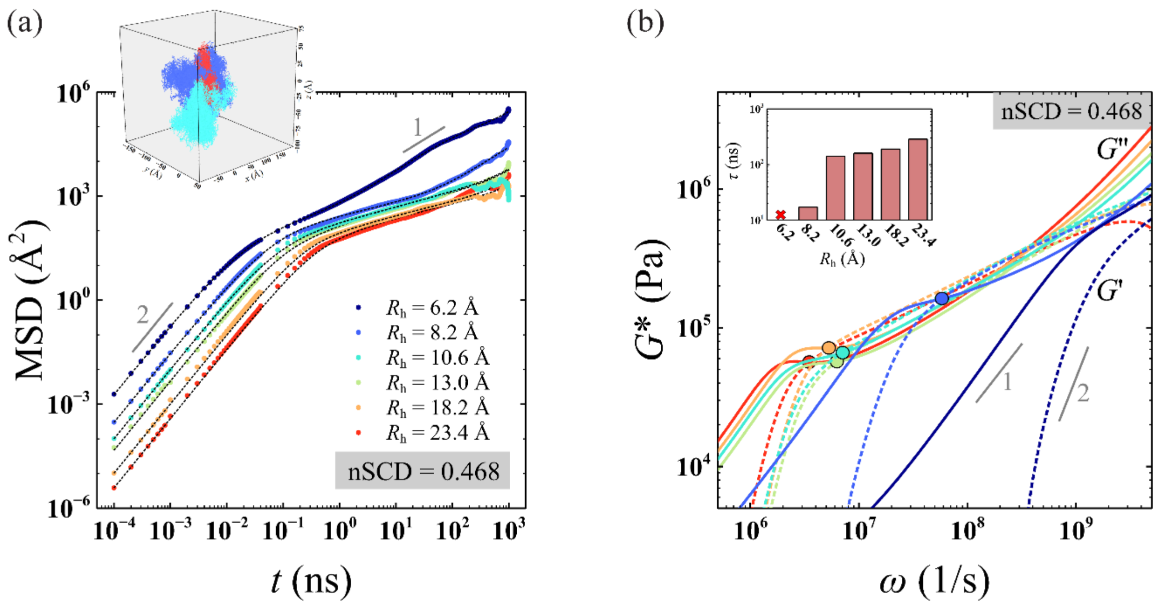


**Fig. S4.** (a) Mean square displacement MSD$(t)$ of the probe particle of different hydrodynamic radius $R_\mathrm{h}$ in the dense phase of the E−K sequence with nSCD = 0.468. The dashed lines are the fits based on the Baumgaertel-Schausberger-Winter-like power law spectrum to the MSD data. The inset shows probe particle's displacement in the three-dimensional Cartesian coordinates for select $R_\mathrm{h}$ values. (b) Elastic $G'$ (dashed line) and viscous $G''$ (solid line) modulus for the dense phase of the E−K sequence with nSCD = 0.468 as a function of $R_\mathrm{h}$, with circles corresponding to the crossover frequency. The inset shows the relaxation time $\tau$ with varying $R_\mathrm{h}$.

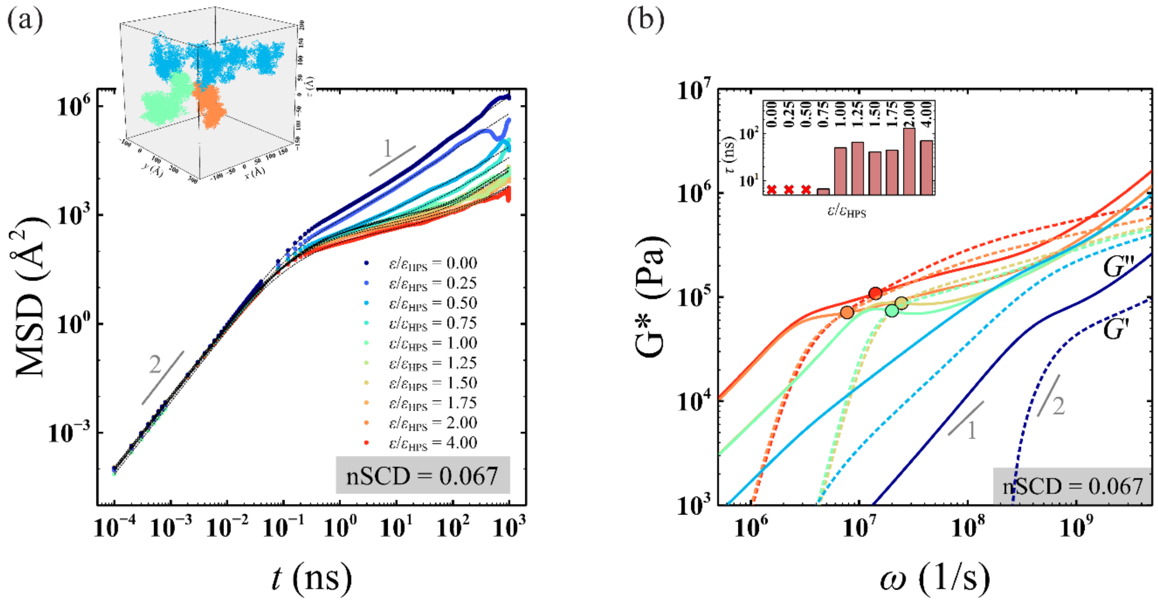


**Fig. S5.** (a) Mean square displacement $\mathrm{MSD}(t)$ of the probe particle for different normalized probe-protein interaction strength $\varepsilon/\varepsilon_{\mathrm{HPS}}$, where $\varepsilon_{\mathrm{HPS}} = 0.20$ kcal/mol, in the dense phase of the E-K sequence with nSCD $= 0.067$. The dashed lines are the fits based on Baumgaertel-Schausberger-Winter-like power law spectrum to the MSD data. The inset shows probe particle's displacement in the three-dimensional Cartesian coordinates for select $\varepsilon/\varepsilon_{\mathrm{HPS}}$ values. (c) Elastic $G'$ (dashed line) and viscous $G''$ (solid line) modulus for the dense phase of the E−K sequence with nSCD $= 0.067$ as a function of $\varepsilon/\varepsilon_{\mathrm{HPS}}$, with circles corresponding to the crossover frequency. The inset shows the relaxation time $\tau$ with varying $\varepsilon/\varepsilon_{\mathrm{HPS}}$.

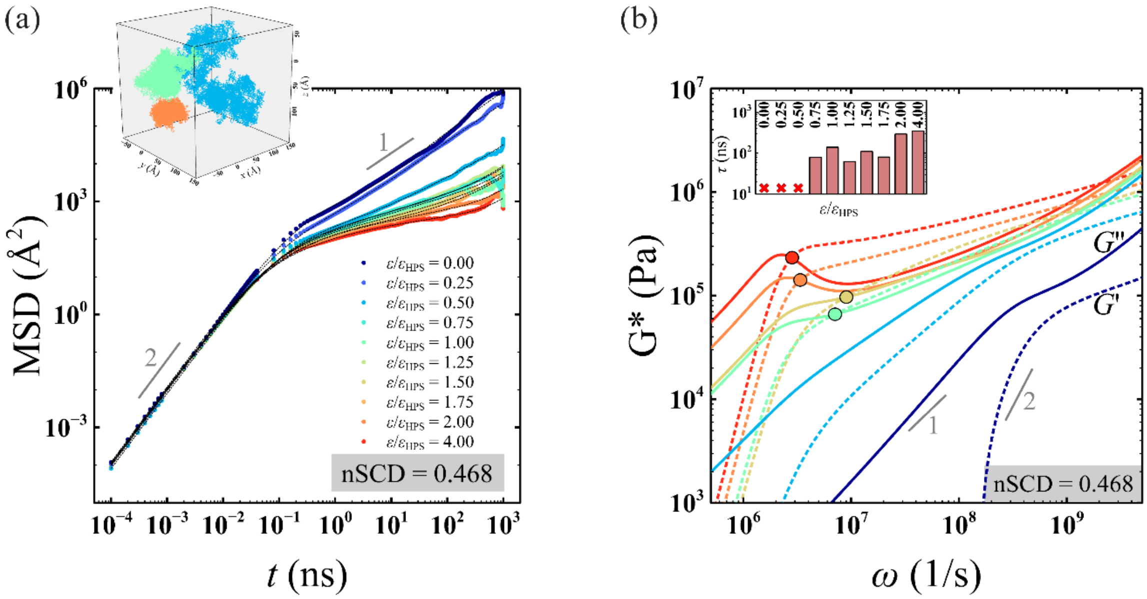


**Fig. S6.** (a) Mean square displacement MSD$(t)$ of the probe particle for different normalized probe-protein interaction strength $\varepsilon/\varepsilon_{\mathrm{HPS}}$, where $\varepsilon_{\mathrm{HPS}} = 0.20$ kcal/mol, in the dense phase of the E-K sequence with nSCD = 0.468. The dashed lines are the fits based on Baumgaertel-Schausberger-Winter-like power law spectrum to the MSD data. The inset shows probe particle's displacement in the three-dimensional Cartesian coordinates for select $\varepsilon/\varepsilon_{\mathrm{HPS}}$ values. (c) Elastic $G'$ (dashed line) and viscous $G''$ (solid line) modulus for the dense phase of the E−K sequence with nSCD = 0.468 as a function of $\varepsilon/\varepsilon_{\mathrm{HPS}}$, with circles corresponding to the crossover frequency. The inset shows the relaxation time $\tau$ with varying $\varepsilon/\varepsilon_{\mathrm{HPS}}$.

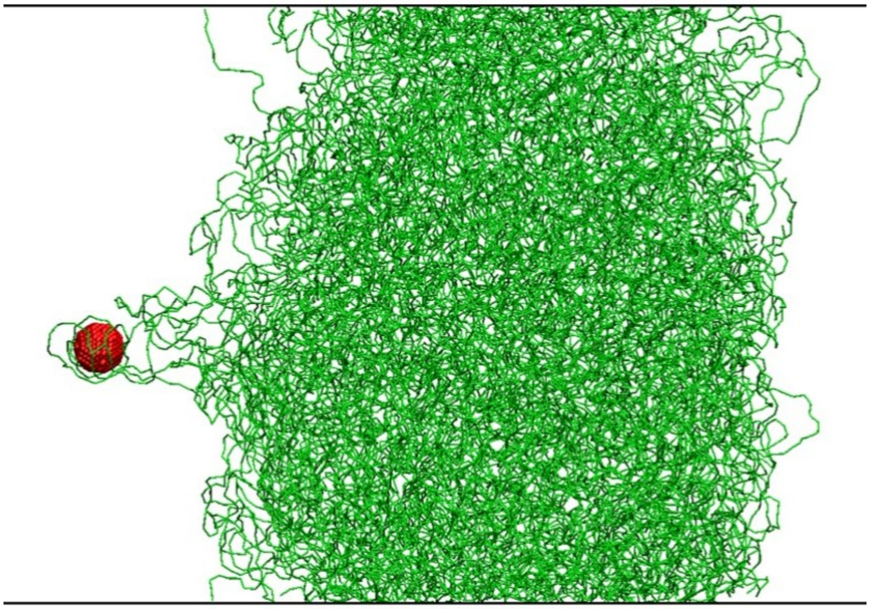

**Fig. S7.** Simulation snapshot showing strong adsorption of E−K protein chains (green) to the probe particle (red) surface, when $\varepsilon/\varepsilon_{\mathrm{HPS}} = 2$, where $\varepsilon_{\mathrm{HPS}} = 0.20$ kcal/mol.

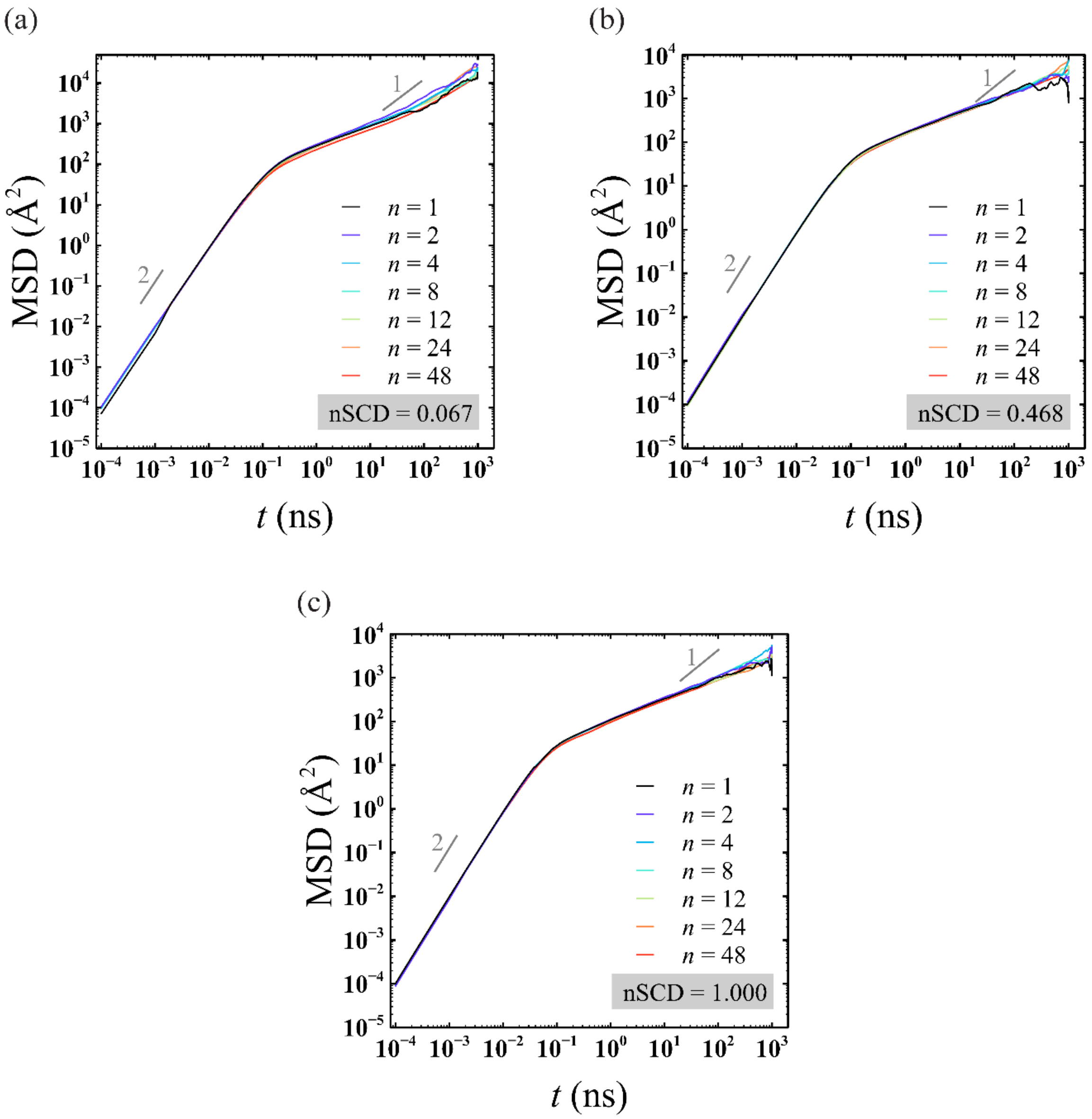


**Fig. S8.** (a) Average mean square displacement MSD($t$) of the probe varying in number $n$ in the dense phase of the E−K sequences for (a) nSCD = 0.067, (b) nSCD = 0.468, and (c) nSCD = 1.000, respectively.

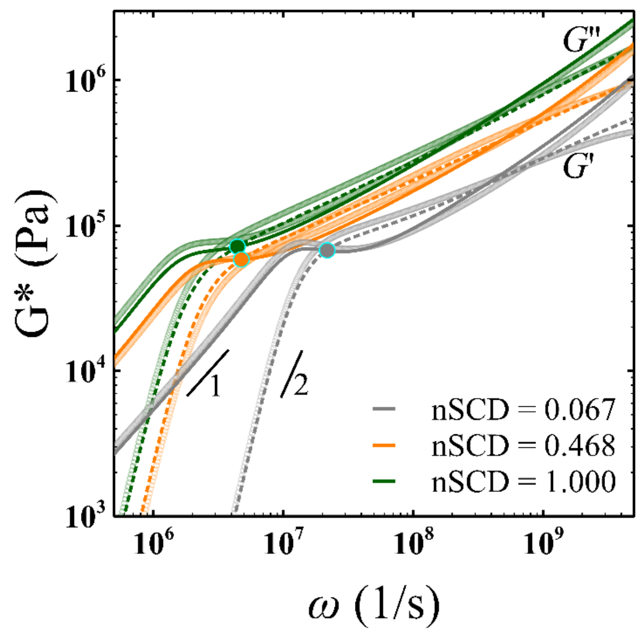


**Fig. S9.** Elastic $G'$ and viscous $G''$ modulus for the E−K sequences with different nSCD. The lines correspond to those obtained based on multiple ($n = 8$) probes and the symbols correspond to those obtained based on a single probe in each dense phase system.

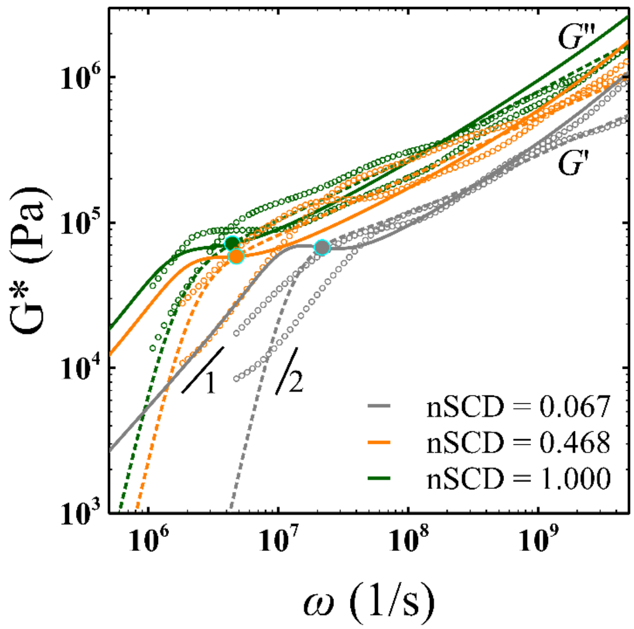


**Fig. S10.** Elastic $G'$ and viscous $G''$ modulus for the E−K sequences with different nSCD. The lines correspond to those obtained based on multiple ($n = 8$) probes from microrheology simulations and the symbols correspond to those obtained from the Fourier transform of shear stress relaxation modulus $G(t)$ computed using the Green-Kubo (GK) relation for each dense phase system.

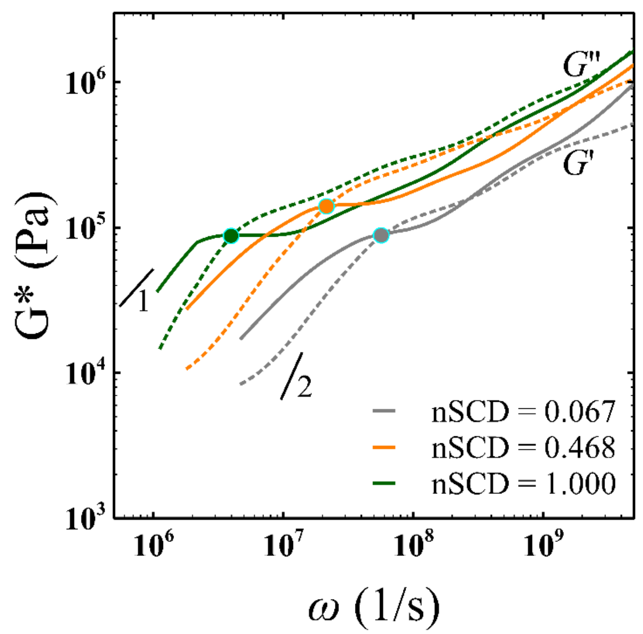


**Fig. S11.** Elastic $G'$ (dashed line) and viscous $G''$ (solid line) modulus for the dense phase of the E−K sequences with different nSCD, obtained from the Fourier transform of shear stress relaxation modulus $G(t)$ computed using the Green-Kubo (GK) relation.

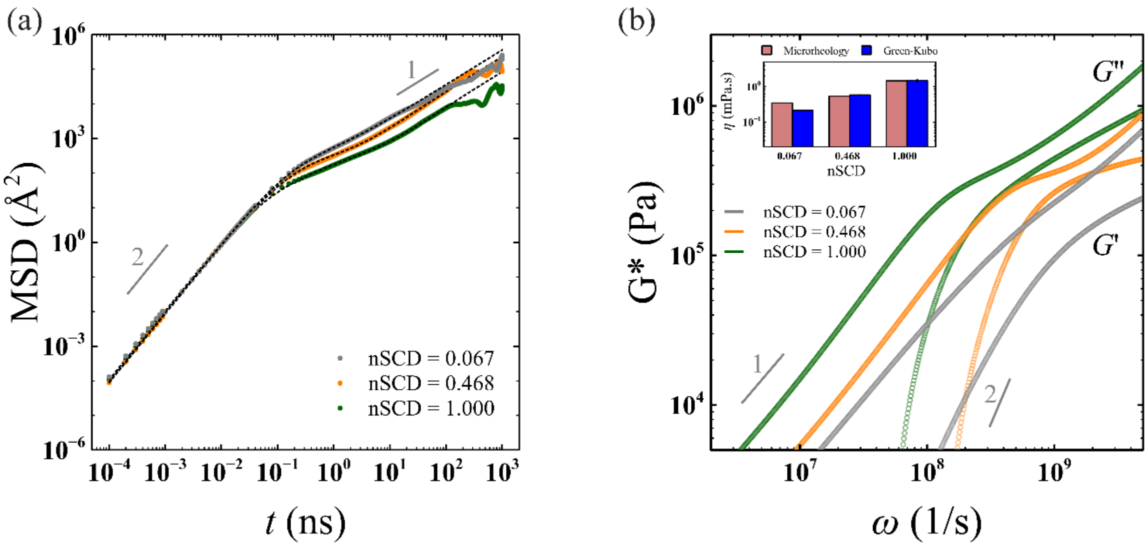


**Fig. S12.** (a) Mean square displacement MSD$(t)$ of the probe particle in the dense phase of different nSCD sequences of $N = 50$. The dashed lines are the fits based on Baumgaertel-Schausberger-Winter-like power law spectrum to the MSD data. (b) Elastic $G'$ and viscous $G''$ modulus for the dense phase of different nSCD sequences of $N = 50$. The insets shows the comparison between the viscosity $\eta$ values obtained from the microrheology simulations and the Green-Kubo method.

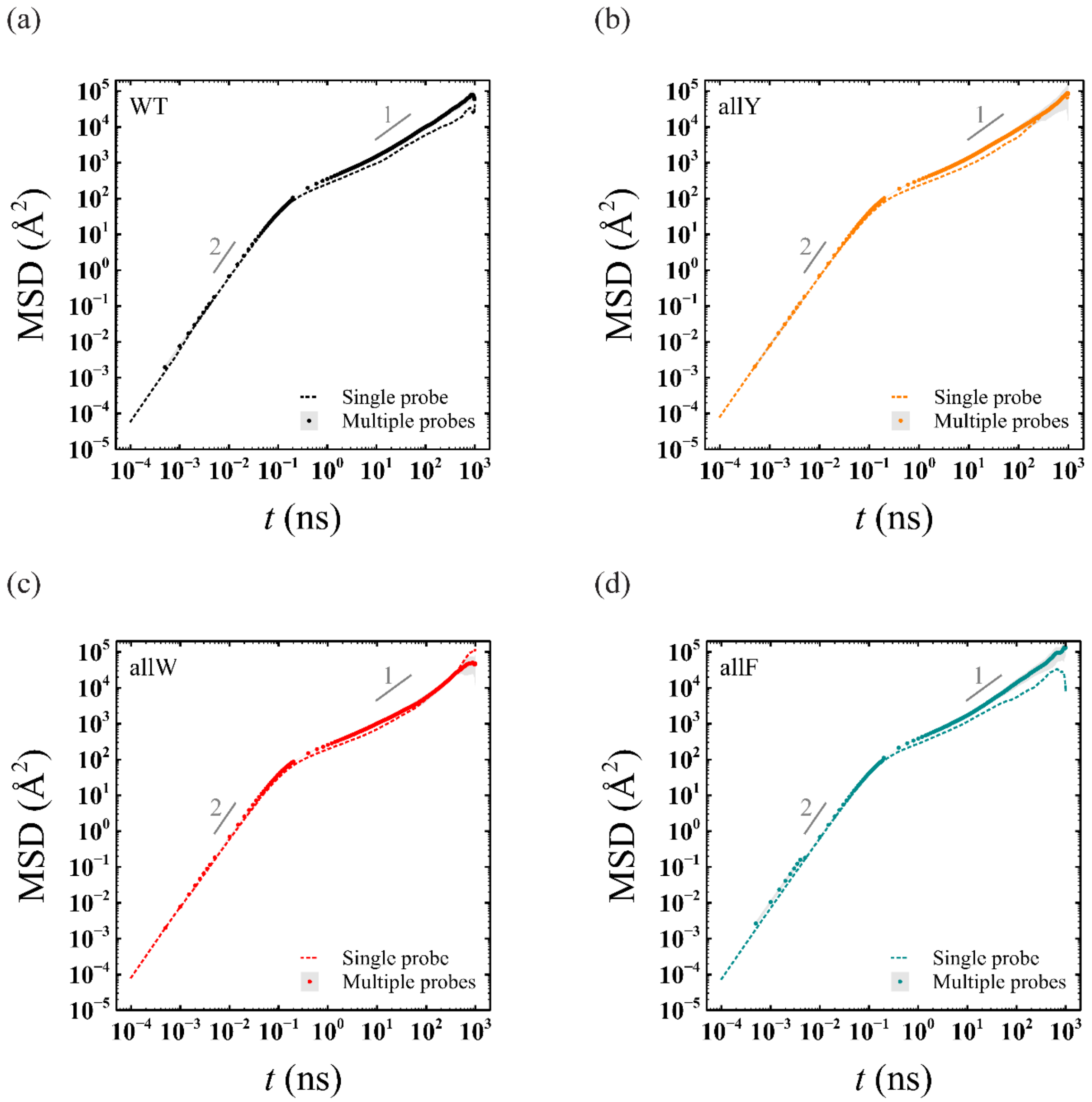


**Fig. S13.** (a) Mean square displacement $\mathrm{MSD}(t)$ of a single probe particle compared with that obtained as an average based on multiple ($n = 6$) probes in the dense phase of the (a) A1-LCD WT sequence and its variants (b) allY, (c) allW, and (d) allF, respectively.

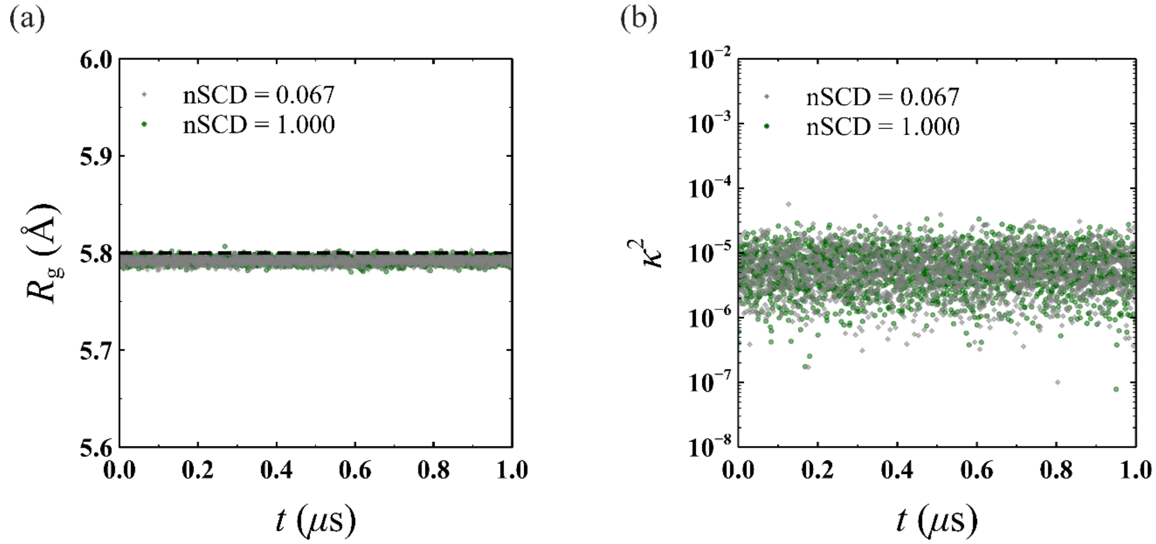


**Fig. S14.** Time evolution of (a) radius of gyration $R_\mathrm{g}$ and (b) relative anisotropy $\kappa^2$ for the probe particle of bare radius $R_\mathrm{b} = 7.5$ Å undergoing Brownian motion in the dense phase of E−K sequences with nSCD = 0.067 and nSCD = 1.000, respectively. The dashed line in (a) corresponds to the expected values for a spherical particle $R_\mathrm{g} = \sqrt{3R_\mathrm{b}^2/5}$.

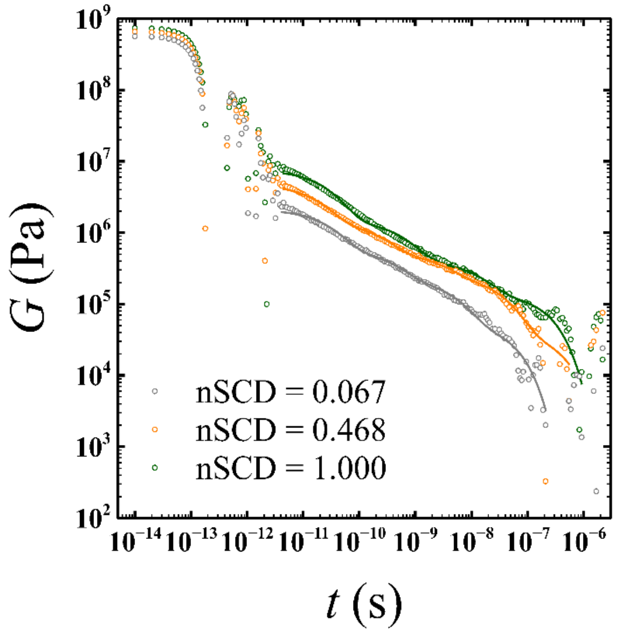


**Fig. S15.** Shear stress relaxation modulus $G(t)$ obtained using the Green-Kubo (GK) relation for the E−K sequence variants with different nSCD. The solid lines at long time scales represent the fit to a series of Maxwell modes.
